# SHINE: SERS-based Hepatotoxicity detection using Inference from Nanoscale Extracellular vesicle content

**DOI:** 10.1101/2025.01.30.635446

**Authors:** Ugur Parlatan, Luke Boudreau, Hulya Torun, Letao Fan, Ugur Aygun, Ayse Aslihan Gokaltun, Demir Akin, O. Berk Usta, Utkan Demirci

## Abstract

Extracellular vesicles (EV) are becoming crucial tools in liquid biopsy, diagnostics, and therapeutic applications, yet their nanoscale characterization remains challenging. In this context, the detection of drug-induced liver injury, *i*.*e*., hepatotoxicity, through extracellular vesicle molecular content remains an unexplored frontier. To this end, we present a label-free surface-enhanced Raman (SERS) spectroscopy approach, which provides rapid EV content analysis under ten minutes and requires only 1.3 microliters of sample. Using hepatic cultures as a model, our platform captures distinct and reproducible EV molecular changes in response to acetaminophen-induced hepatoxicity. Our platform achieves exceptional accuracy with root mean squared error values as low as 3.80%, establishing strong correlations between EV spectra and conventional toxicity biomarkers. Unlike previous EV-SERS studies limited to vesicle identification and disease markers, this approach reveals EV drug-response signatures strongly correlated with conventional toxicity markers. These findings establish EVs as dynamic reporters of cellular drug responses and demonstrate use of SERS-based EV detection of hepatotoxicity.

## Introduction

Extracellular vesicles (EVs) are lipid-bilayer-enclosed biological nanomaterials secreted by virtually all cell types, functioning as nano-messengers that deliver bioactive molecules such as proteins ^1^, lipids^2^, and nucleic acids^3,4^, enabling essential intercellular communication and regulation of biological processes^5^. By encapsulating and transporting molecular signatures that mirror the physiological states of their parent cells, EVs have become central to nano-biomaterial research^6^, drug delivery^7^, medical imaging^8,9^, biomedical diagnostics^10^ and therapeutic development^11,12^. In particular, their molecular cargo can be harnessed for non-invasive monitoring of cellular stress and toxicity, making them ideal candidates for studying drug-induced hepatotoxicity^13–15^.

Drug induced liver injury (DILI), *i*.*e*., hepatotoxicity, remains a leading cause of drug failure during development^16–19^ and withdrawal post approval^20,21^. Standard hepatotoxicity assessment methods, such as liver enzyme assays, histopathological evaluations, and transcriptomic analyses, remain indispensable but are hindered by their invasiveness, lengthy procedures, and limited diagnostic precision^22,23^. Recently, surface-enhanced Raman spectroscopy (SERS) has emerged as a powerful, non-invasive analytical technique^24^ that enhances Raman signals through plasmonic nanostructures^24^. While several studies, including our own^25^ and those of others^26–28^, have explored EV-based SERS sensing for disease diagnostics, the potential of EVs for measuring drug-induced effects, including label-free assessment and rapid classification of cytotoxicity, remains largely untapped.

Here, we present a novel label-free analytical framework, SERS-based Hepatotoxicity detection using Inference from Nanoscale Extracellular vesicle content (SHINE). This framework provides rapid identification and classification of drug-induced hepatotoxicity using a SERS-based platform (**Fig. 1**) by analyzing EVs derived from 2D rat hepatocyte cultures exposed to Acetaminophen (APAP). Specifically, our method utilizes 1.3 µL of sample volume dried on a disposable gold nanopillar-structured SERS surface. Within 10 minutes, we quantified APAP-induced hepatotoxicity by capturing the molecular changes in EV cargo two days after drug exposure. Our regression model, trained on SERS spectral features, successfully predicted APAP doses across independent cultures with root-mean-square error (RMSE) values as low as 1.50 mM. This study demonstrates the potential of EV-SERS integration as a precise, non-invasive, and scalable approach for hepatotoxicity screening, offering a new avenue for improving preclinical drug development and toxicity prediction.

**Figure 1:**
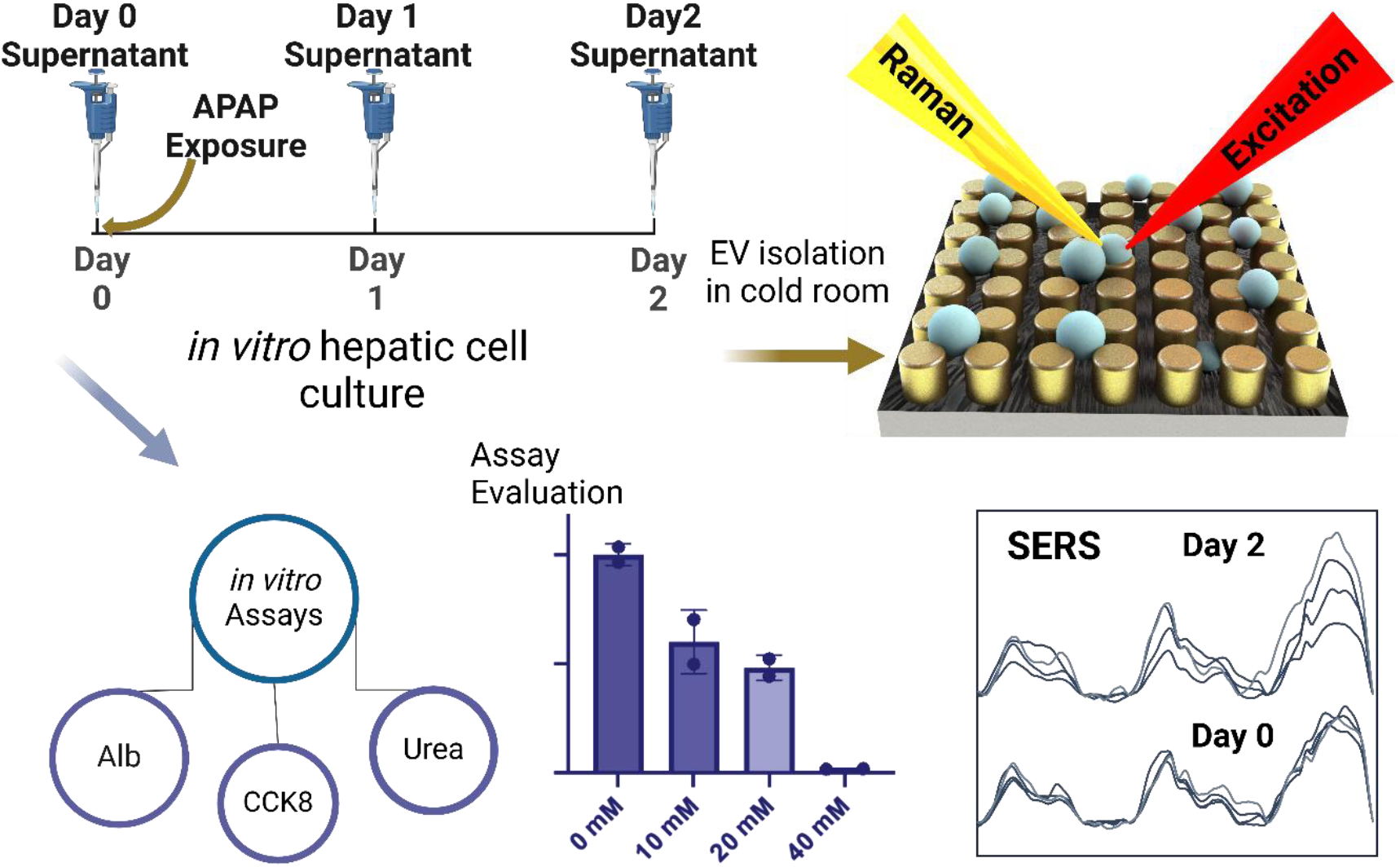
Experimental approach used in this study. Hepatic cell cultures were prepared in vitro on plates for various doses and durations. Extracellular vesicles were isolated from the supernatants of these cultures and prepared on gold nanopillar SERS substrates. SERS spectra were compared across different days and doses. Concurrently, functional assay measurements were taken to assess cell viability as the dose increased. These viability results were then compared with the SERS intensities (Created in https://BioRender.com).

## Results

We used a SERS-based, label-free approach, to evaluate the changes in EV content secreted by rat primary hepatocytes under different APAP exposures, representing various levels of hepatotoxicity. Increasing doses of APAP led to a dose-dependent decline in hepatocyte viability, functionality, and morphological integrity over 2 days (**Fig. 2**). Cell viability measurements on day 2, based on a CCK8 assay, indicated an IC50 of 14.55 mM (**Fig. 2A**). Concurrent functional assays on day 2 showed significant reductions in albumin secretion (IC50: 12.6 mM, **Fig. 2B**) and urea excretion (**Fig. 2C**) with increasing APAP doses. Morphological analysis on day 2 corroborated these findings, revealing extensive cellular damage and culture disintegration at 40 mM (**Supplementary Figure 1**). These results collectively underscore the severe cytotoxic effects of APAP at high doses on primary rat hepatocytes.

**Figure 2:**
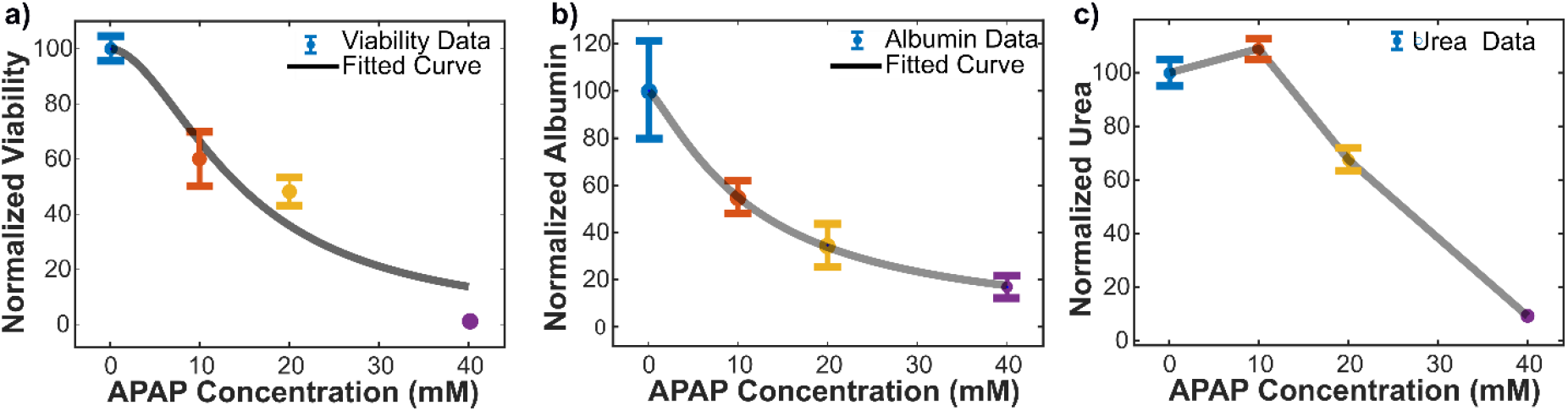
Viability, functionality, and morphology of primary rat hepatocyte cultures with varying doses of Acetaminophen exposure. **A)** The normalized viability of cell samples via CCK8 assay at different concentrations of acetaminophen (APAP) at the end of the experiment (after 2 days of incubation with APAP). The 2-day IC_50_ was determined to be 14.55 mM. **B)** The normalized change of albumin concentration in cell culture medium at the endpoint of the experiment. The day 2 IC_50_ was determined to be 12.6 mM. **C)** The normalized change in urea concentration in cell culture medium at the endpoint of the experiment.

Next, we isolated EVs from hepatic culture supernatants using our ExoTIC platform^29^ as described in the methods section. NTA analysis revealed that the EVs primarily fall within the small EV size range (50 – 220 nm) (**Fig. 3A**). As seen in **Fig. 3A**, the Day 2 – 40 mM measurements have a large variance in concentration since the levels are close to limit of detection of the device^30^. We, thus, used a complementary technique, iSCAT, and confirmed the presence of EVs in the low concentration range and quantified the decrease in the concentration in day 2 for 40 mM APAP (**Fig. 3B**). The tables summarizing the size profiles from NTA and iSCAT can be seen in **Supplementary tables 1 and 2**. We confirmed the identity of isolated EV samples using flow cytometry with a CD63 antibody, a tetraspanin marker. High CD63 positivity (>90%), in our samples, indicates that most analyzed particles are EVs (**Fig. 3C)**. Extended version of the flow cytometry data can be seen in **Supplementary Figure 2. Figure 3D** shows a Western blot analysis of EV-specific markers CD63, TSG101, and Flotillin-1 (Flot1), which are strongly expressed, supporting the presence of EVs. In addition to CD 63, TSG101, an ESCRT complex protein, and Flotillin-1, a membrane-associated lipid raft protein, further confirm the EV identity. The absence of a Calnexin band, an endoplasmic reticulum marker, verifies the purity of the EV preparation by excluding cellular contamination. TEM further validated the morphology and size of the hepatic EVs (**Fig. 3E**).

**Figure 3:**
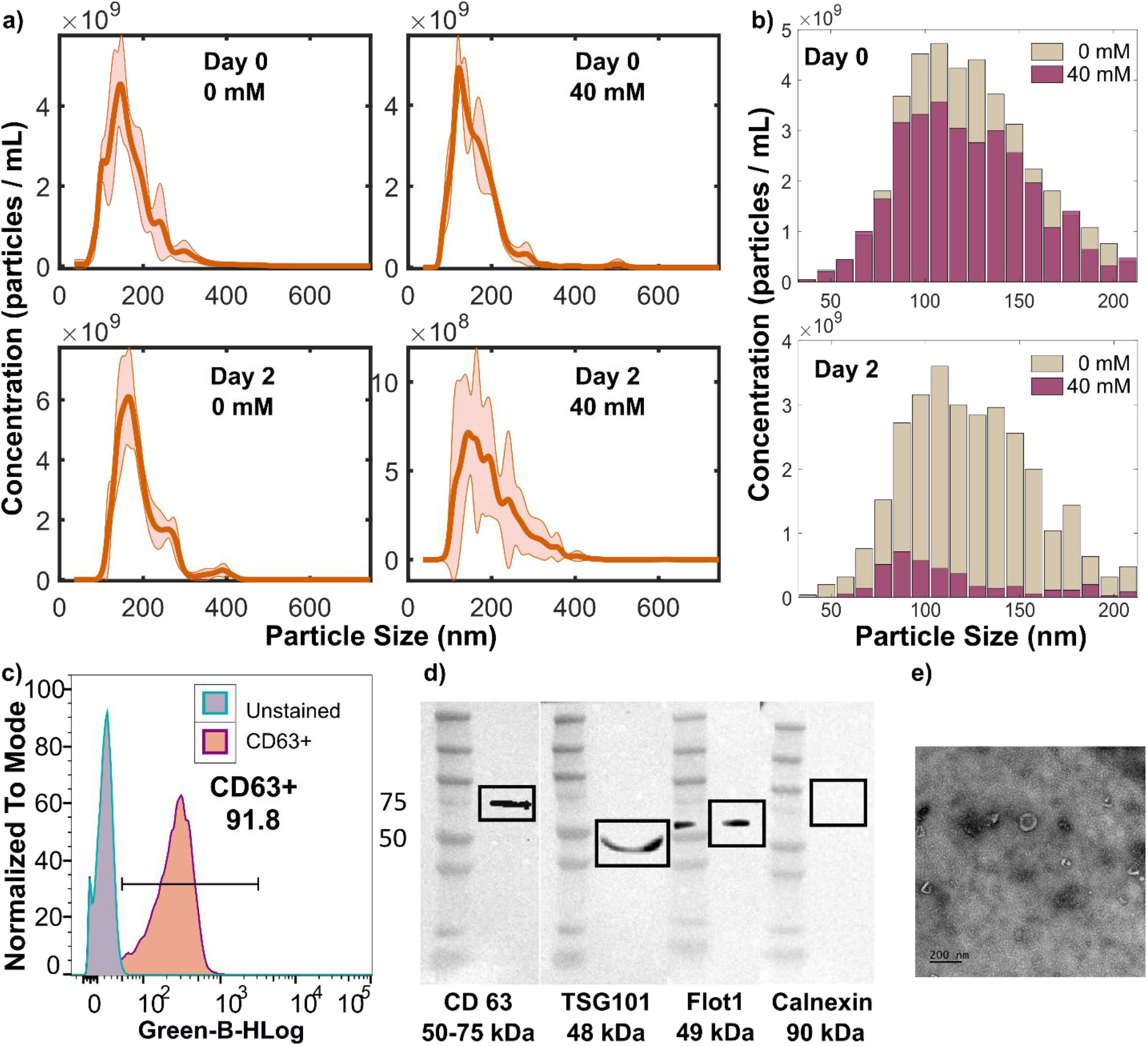
Characterization hepatic EVs. **A)** Nanoparticle tracking analysis (NTA) of EV size and concentration for Day 0 and Day 2 samples under 0 mM and 40 mM APAP exposure. Shaded areas represent standard errors. **B)** iSCAT analysis comparing EVs under 0 mM and 40 mM APAP exposure. Top: Day 0 samples; Bottom: Day 2 samples. **C)** Flow cytometry was utilized with and without CD63 antibodies to prove the presence of small EVs from Day 0 samples. **D)** Western blot analysis of EV markers CD63, TSG101, and Flotillin-1, with no detectable signal for Calnexin, confirming the absence of cellular contamination in the EV preparations. **E)** Transmission electron microscopy (TEM) was used to show the particles have the expected small EV shapes within sizes smaller than 220.

We then evaluated dose-dependent effects of APAP exposure using label-free molecular vibrational analysis. Specifically, we used SERS data on EVs derived from culture supernatants derived from 0, 10, 20, and 40 mM APAP exposure across two independent sets (Set 1 and Set 2) to assess hepatotoxicity. To this end, we applied principal component analysis (PCA) and trained regression models to explore the spectral differences. **Figures 4A and 4B** present the average normalized spectra, highlighting reduced intensities with increasing drug concentration on Day 2 particularly in the 810–1600 cm^-1^ range, while we observed no significant differences before dosing on Day 0, indicating dose-dependent spectral changes.

**Figure 4:**
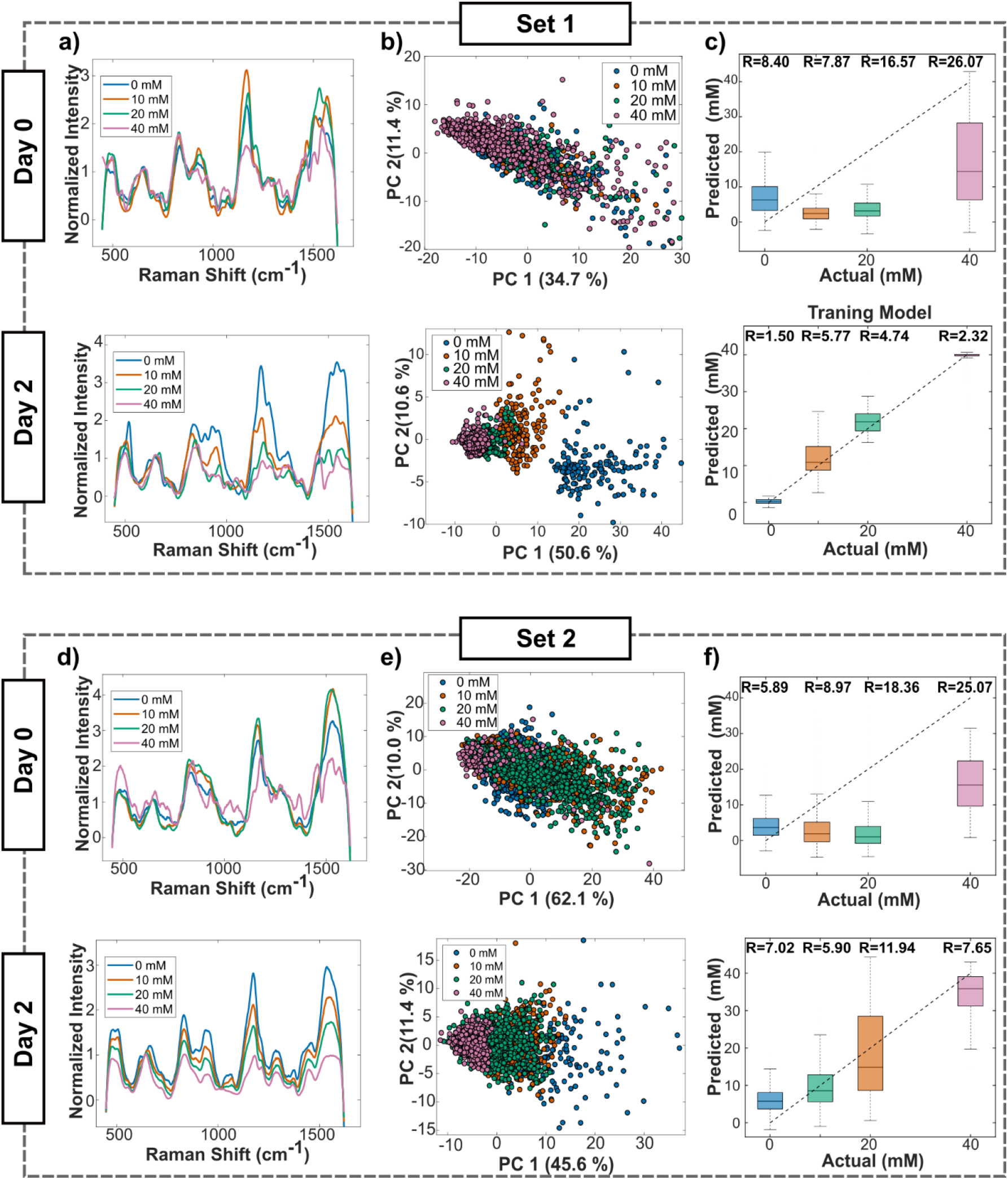
SERS analysis of EVs from hepatic cell cultures. A,D) Dose-response spectra for days 0 and 2 (Sets 1 and 2). B,E) PCA score maps for dose responses on days 0 and 2 (Sets 1 and 2). C) Gaussian Process Regression (GPR) model calibrated with 80% of Set 1 day 2 data. Top: Predictions for Set 1 day 0. Bottom: Test set predictions for Set 1 day 2. F) Predictions for Set 2 day 0 (Top) and day 2 (Bottom) using the Set 1 day 2 model.

To assess whether spectral analysis of EV content can distinguish differences in APAP exposure, we performed principal component analysis (PCA) (**Figs. 4C and 4D)**. On Day 2, PCA revealed well-separated clusters corresponding to concentration levels in both sets. In contrast, day 0 data showed no visible clustering (See **Table S1** for overlap analysis). This demonstrated that spectral differences induced by hepatotoxicity became more pronounced on Day 2, highlighting the progressive effect of hepatotoxicity and the improved ability to distinguish concentrations over time.

Next, we created a regression model using the Set1 – Day 2 dataset by applying Gaussian Process Regression (GPR)^31^ analysis (**Figs. 4E and 4F)**. The GPR model demonstrated strong predictive performance on the test set for Set 2 – Day 2 dataset, with good agreement between actual and predicted concentrations, as reflected by RMSE values. To validate the model’s prediction ability, we analyzed Day 0 data as a negative control, where the model consistently failed to predict concentrations accurately across both sets, showing substantial deviations regardless of concentration levels.

We also sought to identify SERS wavenumbers that show the highest correlation with the traditional markers of hepatic cell viability across the APAP dose range. Specifically, we calculated the Pearson correlation between the normalized SERS intensities for each wavenumber with CCK-8, albumin and urea measurements and rank-ordered them. **Figure 5**, where in each panel the x-axis represents the SERS wavenumber intensity and y-axis represents the normalized valued of different traditional markers, illustrates the correlation of select SERS wavenumbers which showed the highest correlations (>0.95 with albumin) and are biochemically relevant. The six highly correlated SERS intensities were around 739 cm^-1^ (**Fig. 5A**), 959.8 cm^-1^ (**Fig. 5B**), 1249.8 cm^-1^ (**Fig. 5C**), 1525.3 cm^-1^ (**Fig. 5D**), 1576.2 cm^-1^ (**Fig. 5E**), and 1602.1 cm^-1^ (**Fig. 5F**) which showed high linearity against both the CCK-8 and albumin signals. A detailed matrix of all correlations among the SERS and traditional marker data is tabulated in **Table 1**. We note that albumin and CCK-8 measurements are highly correlated with each other and albumin is considered one of the most reliable viability markers for hepatic cultures^32^. Urea measurements, on the other hand lag these two both temporally and in a dose dependent manner ^33^, which is also evident in our results (**Fig. 2 and Table 1**). **Table 2** presents the tentative peak assignments for the selected SERS wavenumbers and includes the corresponding molecular vibrational bonds and the most probable macromolecular associations retrieved from the literature.

**Figure 5:**
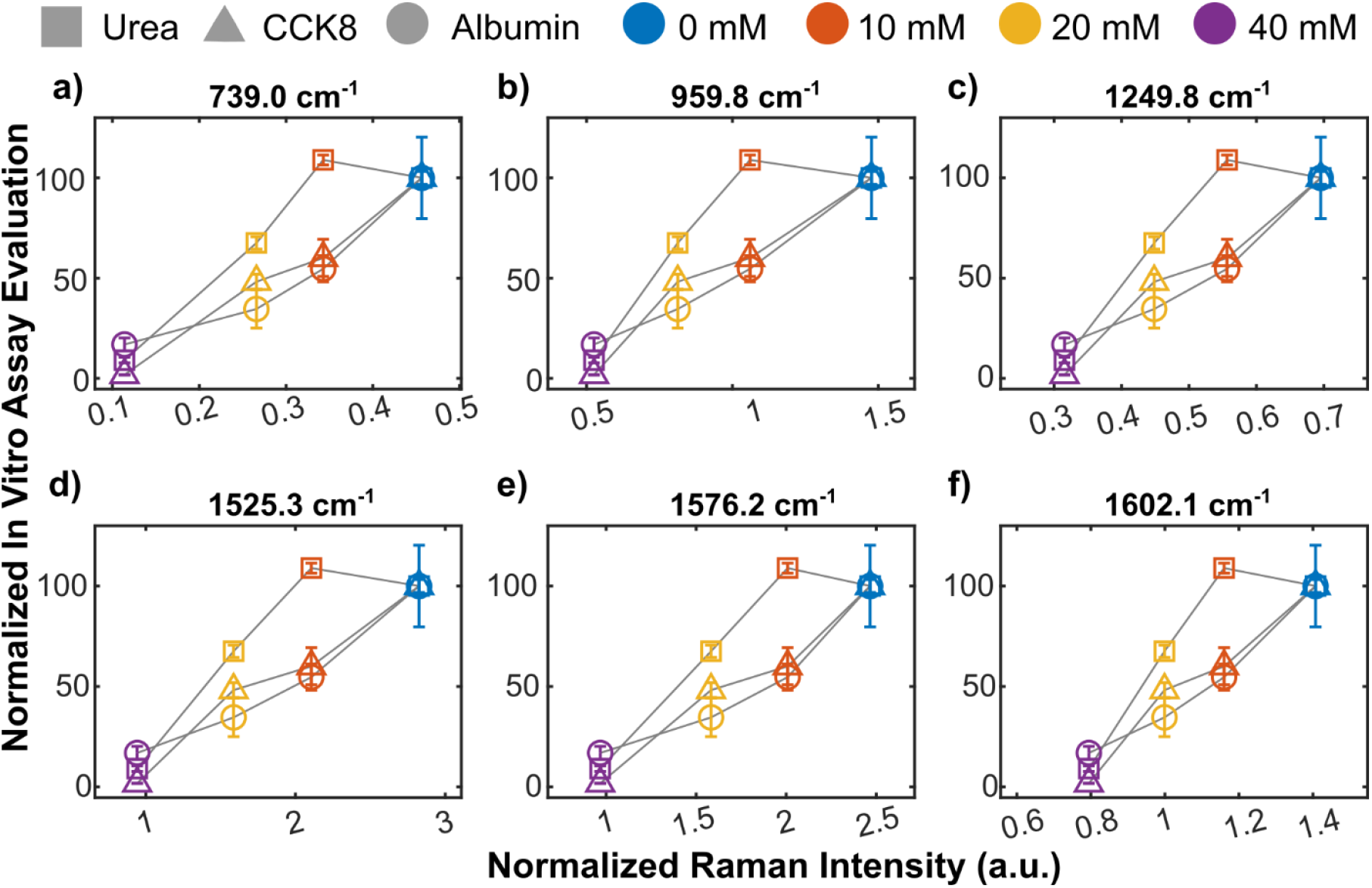
SERS intensity vs viability test results using functional assays on albumin, CCK8 and urea. The relations of these analytes with SERS peak intensities at **A)** 739, **B)** 960, **C)** 1250, **D)** 1525, **E)** 1576, **F)** 1602 cm^-1^ are displayed in the figure.

**Table 1:**
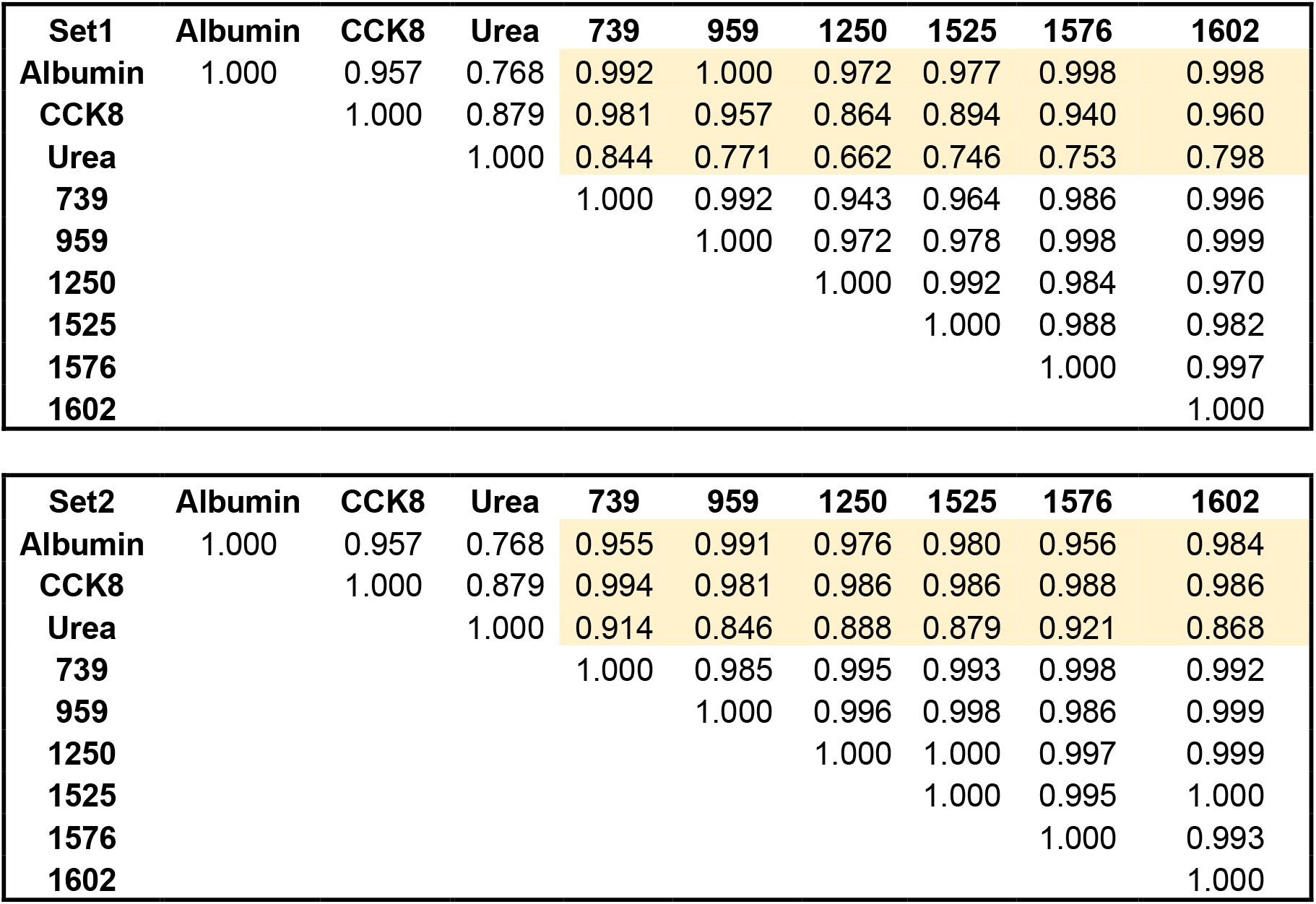
Correlation coefficients between normalized viability scores via albumin, CCK8 and urea and SERS intensities at 739, 960, 1250, 1525, 1576 and 1602 cm^-1^.

**Table 2:** Tentative band assignments of the selected Raman bands.

| Wavenumber | Bond Vibration | Assignment | Reference |
| --- | --- | --- | --- |
| 736 | Pyridine ring breathing | RNA, observed in miR-122 | 43 |
| 960 | $\nu(\text{PO}_4^{3-})$ | Ribose | 34 |
| 1185 | Ring $\delta_{\text{ip}}(\text{C-H})$ | Saccharides, lipids, nucleic acids | 35,36 |
| 1250 | Amide III | Proteins | 37 |
| 1525 | $\nu_{\text{ip}}(\text{C=C})$ | C, G, RNA assignment | 38,39 |
| 1576 | Ring breathing modes | A,G | 40 |
| 1602 | Pyridine ring stretching | RNA, observed in miR-122 | 41,42 |

## Discussion

In this study, we demonstrated the potential of SHINE, a SERS-based EV analysis platform to detect drug-induced liver injury, through dose-dependent molecular signatures. We observed a consistent decrease in SERS intensities between 810-1600 cm^-1^ as we increase the APAP dosage. Using this dataset and the Gaussian Process Regression (GPR) model, we achieved RMSE values as low as 1.50 mM (3.8%) across two independent datasets, indicating that the model has the ability to accurately predict an the varying APAP dosages via SERS intensities. A correlation analysis between SERS intensities and *in vitro* markers revealed key wavenumbers, such as 739, 960, 1250, 1525, 1576 and 1602 cm^-1^, which correlate strongly (r > 0.94) with both albumin and CCK8 measurements across all APAP doses. As indicated in **Table 2**, these wavenumbers align with known nucleic acid signatures^34–42^ most predominantly with those of RNA. This result indicates regulation of RNA and its packaging into EVs under varying APAP exposure. This finding is supported by the previous studies which have identified cellular and EV-derived miRNAs, such as miR-122^43^, miR-192^44^, miR-193^45^, as key markers of drug induced liver injury.

Our study has several limitations. First, due to the high dilution (1:10,000) of the samples, we did not obtain measurable signal from the EVs in certain measurements. Thus, we utilized a cluster analysis to filter out the unknown spectral signature. Second, surface-based sensing methods depend on surface stability, uniformity, and repeatability and may suffer from batch-to-batch variability. To control this, we ensured signal integrity by performing all measurements using surfaces from a single batch (See **Supplementary Figure 3A** for surface uniformity). Third, due to the low number of APAP dosage used, we observed slight deviations from linearity in our regression model when applied to independent validation set. Despite this limitation, we still achieved RMSE values down to 5.90 mM (14.8%) for this validation set. Further improvement in such regression can be achieved by expanding the number of dosage and increasing number of both the training and the validation sets. Finally, in the absence of a normalization factor based on the number of EVs analyzed, we employed a spectral normalization approach using a single peak intensity.

The significance of this study lies in developing an EV-based model that quantifies hepatotoxicity through molecular signatures detected by SERS. This approach enables real-time, non-invasive monitoring of drug-induced liver toxicity, advancing the predictive capacity of preclinical drug assessments. Future research could explore dynamic EV responses across a broader range of hepatotoxic compounds and their associations with various miRNA signatures. Expanding the dataset by incorporating additional dosages and donors could further improve the model’s generalizability, positioning the SHINE platform as a standard tool for non-invasive drug toxicity assessment and preclinical screening.

## Methods

### Cell Culture and Analysis Materials

Two different cell culture media were used during culture. First, primary rat hepatocyte culture media (C + H) was prepared with high glucose (4.5 g/L) Dulbecco’s modified eagle’s medium (DMEM; Life Technologies, CA, United States) and was supplemented with 10% fetal bovine serum (FBS, Sigma, St. Louis, MO, United States), 2% penicillin-streptomycin, 7.5 μg/mL hydrocortisone, 20 ng/mL epidermal growth factor (EGF) and 14 ng/mL glucagon, was used for seeding and initial overnight maintenance. Hepatocyte Culture media, as defined for the rest of the protocol, is the HCM Hepatocyte Culture Medium BulletKit including HBM Basal Medium (CC-3199) and HCM Singlequot Supplements (CC-4182) (0.5 mL transferrin, 0.5 mL ascorbic acid, 0.5 mL HEGF, 0.5 mL insulin, 0.5 mL hydrocortisone, 10.5 mL BSA (Fatty Acid Free), and 0.5 mL GA-100). HCM Hepatocyte Culture Medium BulletKit including HBM Basal Medium (CC-3199) and HCM Singlequot Supplements (CC-4182) were purchased from Lonza Bioscience (Walkersville, MD). Hepatocyte Culture media was used after the first day of culture and for the duration of the experiment.

12 well tissue culture plates were purchased from CellTreat Scientific Products (Pepperell, MA, United States). GenClone 25-508 Dulbecco’s phosphate-buffered saline (PBS) was purchased from Genesee Scientific (Research Triangle Park, NC, United States). Trypan blue was purchased from Sigma Aldrich, United States. Geltrex™ LDEV-Free Reduced Growth Factor Basement Membrane Matrix (A1413202), and 12 and 96 well plates were purchased from Invitrogen by Thermo Fisher Scientific (Carlsbad, CA, United States). Cell counting kit-8 (CCK8) was purchased from ApexBio (ab228554; Boston, MA, United States). The urea assay kit was purchased from Stanbio Laboratory (0580-250). Tween 20 (P2287), albumin from rat serum (A6414), o-phenylenediamine dihydrochloride (OPD) tablets (10 mg substrate per tablet) (P8287), hydrogen peroxide (216763), sulfuric acid (320501), dimethyl sulfoxide (DMSO) (472301), and acetaminophen (A7085) were purchased from Sigma Aldrich (St. Louis, MO, United States). Sheep anti-rat albumin HRP conjµgated (A110-134P) (1 mL at 1 mg/mL) was purchased from Fortis Life Sciences (Waltham, MA, United States).

### Primary Rat Hepatocyte (PRH) Isolation

Primary rat hepatocytes (PRH) were freshly isolated from 10 to 12 weeks-old adult female (180– 200 g) Lewis rats (Charles River Laboratories, United States). The Cell Resource Core (CRC) performed the isolation according to protocol #2011N000111, as approved by the Institutional Animal Care and Use Committee (IACUC) at Massachusetts General Hospital (MGH). After complete digestion of the liver, a yield of 400–600 million primary rat hepatocytes with greater than 85% viability was given for use. Cell count and viability were provided by quantification with a hemocytometer count after trypan blue staining.

### 2D Rat Hepatocyte Monolayer Culture and Acetaminophen Dosage

Prior to cell seeding, 12-well tissue culture plates were coated with 500 μL 1% Geltrex in PBS, and incubated at 37°C in a humidified environment containing 5% CO_2_. Next, the remaining solution was aspirated and the well was washed with PBS. The PRH cell suspension was diluted to 560,000 viable cells/mL in C+H medium and 1 mL was added to each well. After introduction of the hepatocyte suspension to the wells, the plate was shaken gently in an alternating horizontal and vertical pattern to uniformly distribute the cells within the wells. The plates were then incubated for 45 minutes at 37°C with 5% CO_2_. Following the initial incubation, the medium was aspirated and each well washed with warm PBS to remove dead or non-adherent cells. Finally, 1 mL of C+H medium was added. After 24 hours, the media was aspirated and cells were washed with PBS. For the second 24 hours of culture, 1 mL of Hepatocyte Culture medium with 2% Geltrex by volume (2% Geltrex-media) was added to each well.

The 24 hours after the introduction of 2% Geltrex-media, media from each well was collected and stored at -80°C. These samples were later unfrozen for analysis and divided by volume based on experimental usage. The 1 mL samples were split into two parts, a 600 μL aliquot for exosome analysis and 400 μL for functionality assay completion. Following collection of media, a CCK8 assay was completed.

Following absorbance readings, the media from each well was aspirated and washed with PBS. The well plate was then split into four groups and introduced to HCM-2% Geltrex media with either control (no added acetaminophen), 10mM, 20 mM, or 40 mM acetaminophen, diluted from a 2M stock of acetaminophen in DMSO. 24 hours and 48 hours after initial dosage (Day 1 and Day 2), the collection and CCK8 dosage was repeated, and the cells were dosed with acetaminophen-media again.

### CCK8 Assay and Brightfield Imaging

After collection, each well was washed with PBS before 500 µL of Hepatocyte Culture media was added for use in the CCK-8 viability assay. 50 µL of CCK-8 were then added to 8 of the 12 wells and incubated for 1 hour at 37°C with 5% CO_2_. After incubation, triplicate samples of 80 μL were pipetted into a 96-well plate and their absorbance values were read on a microplate reader (Molecular Devices, SpectraMax® ID3) at 450 nm.

Following CCK8 assay and dosing with acetaminophen-media each day, each well was imaged using an EVOS™ M5000 microscope. Morphological changes were used to qualitatively validate the other viability and functional assay results.

### Albumin Quantification

Albumin secretion in PRH cultures was determined using an enzyme-linked immunosorbent assay (ELISA) protocol. The following solutions were prepared prior to beginning the assay: substrate buffer (5.1 g citric acid and 7.29 g sodium phosphate dibasic in 1 L distilled water), 8N sulfuric acid (22.2 mL concentrated sulfuric acid in 77.8 mL DI water), and PBS-Tween 20 solution (0.05 v/v% Tween 20 in PBS).

96 well plates were coated with 100 µL of rat albumin at a concentration of 50 µg/mL. The plate was then covered with an adhesive plate sealer, and incubated overnight at 4°C. After incubation, plates were washed four times with PBS-Tween solution. After washing, 50 µL of sample or standard were added in triplicate. Standards were created by serially diluting 100 µg/mL albumin and added to each individual plate. Following introduction of samples/standards, 50 µL of antibody solution, stock antibodies diluted 1:10000 in PBS-Tween, was added to each well. The plate was then covered with an adhesive plate sealer and incubated for 90 minutes at 37°C.

At the end of the incubation, the plate was washed again with PBS-Tween solution four times. Subsequently, 100 µL of OPD development solution, a mixture of one pill of OPD, 25 mL substrate buffer, and 10 µL of 30% H2O2, was introduced via multichannel pipette at set intervals. After 5 minutes, the reaction was quenched with the addition of 50 µL of 8N sulfuric acid at the same interval as previously before measuring the difference between 490 and 650 nm on a microplate reader (Molecular Devices, SpectraMax® ID3).

### Urea Quantification

Urea excretion of PRH cultures was quantified using a Stanbio urea BUN assay kit following the manufacturer’s protocol. The urea assay reagent was prepared by mixing one part BUN color reagent and two parts BUN acid reagent. Next, 10 μL of either standard or sample are pipetted in duplicate into a 96 well plate, and combined with 150 μL of urea assay reagent. The plate was then covered with a plate sealer and incubated for 90 minutes at 60°C. After removal from the incubator, the plate was cooled at room temperature for 10 minutes. The plate was finally read on a microplate reader (Molecular Devices, SpectraMax® ID3) at 540 nm.

### Extracellular Vesicle Isolation

All EV isolation steps were performed in a cold room to prevent gelation, which could reduce the number of samples and lead to larger size distributions due to aggregation. Supernatants from triplicate samples were pooled to increase the yield of EVs from each sample. The total volume was brought to 5 mL by diluting with PBS. The diluted sample was then processed using our ExoTIC platform^29^ to select nanoparticles within the 50 to 220 nm size range. After the initial isolation, a second wash step was conducted with 5 mL of PBS. This platform is now well established for yield and intact EV isolation outcomes in various applications including cancers ^46–48^, cardiovascular diseases^49–51^, and immunology ^52^.

### Extracellular Vesicle Characterization

We characterized EVs using transmission electron microscopy (TEM), nanoparticle tracking analysis (NTA), flow cytometry, interferometric scattering microscopy (iSCAT) and western blotting. For TEM, 10 µl EVs were loaded onto the grid (Electron Microscopy Sciences – EM Sciences FCF-300-CU), and incubated with 1% uranyl acetate before imaging with a JEOL JEM 1400 120 kV TEM microscope (JEOL Inc., USA) equipped with a Ultrascan digital high-resolution camera (Gatan Inc., Pleasanton, CA). NTA NanoSight equipped with 430 nm laser (NS300, Malvern Panalytical, UK) quantified and profiled EVs in terms size and concentrations. Bead-assisted flow cytometry was performed to further characterize EVs, using 4% w/v, 4 µm aldehyde-sulfate latex beads (A37304, ThermoFisher, Carlsbad, CA, United States), and staining with the EV tetraspanin marker CD63 (#556019, BD Pharmingen, San Jose, CA, USA) antibody. Western blot analysis of EVs was performed using Calnexin (#PA1-30197, Invitrogen, Carlsbad, CA, United States), Flotillin-1 (#A6220, Abclonal, Woburn, MA, USA), TSG101 4A10 (#NB200-112, Novus Bio, Centennial, CO, USA) and CD63 (#556019, BD Pharmingen, San Jose, CA, USA) antibodies, according to the insights provided in MISEV 2023 guidelines ^53^.

### Interferometric Scattering Microscopy

To analyze the concentration and size distribution of Extracellular Vesicles (EVs), a custom-built interferometric scattering (iSCAT) microscope is employed^54,55^. iSCAT imaging enables the visualization of nanoscale particles by leveraging the interference between the scattered light from the nanoparticles and the background light reflected from the sample substrate. This interference generates a signal on the camera plane that correlates with the particle size. In this study, EVs are immobilized on a silicon (Si) substrate coated with a 100 nm-thick SiO_2_ thin film.

The substrate is illuminated in a widefield configuration using a 530 nm LED light source (M530L4, Thorlabs, Newton, NJ, USA), which is focused onto the back focal plane of a 50x objective lens (LMPlanFL, Nikon Corporation, Tokyo, Japan). The reflected and scattered light are collected through the same objective lens and subsequently imaged onto a CMOS camera (acA2440, Basler AG, Ahrensburg, Germany). For each measurement, a sample spot of 0.2 µL is deposited using a standard pipette and allowed to dry for 15 minutes to immobilize the particles.

The sample spot is then scanned, and z-stacks of interferometric images are acquired using a motorized sample stage. The best-focused images are selected based on the particle intensity in the z-stack. Particle detection and counting are performed using a custom MATLAB script. The total particle count in the interferometric images is converted into a concentration (particles/mL) by accounting for the sample volume.

### Surface Enhanced Raman Spectroscopy

Surface-enhanced Raman spectroscopy (SERS) experiments were conducted using a custom-built setup with a 785 nm, 100 mW diode laser (CrystaLaser, Reno, NV, USA). The laser output power was reduced to 5 mW using a half-wave plate and a polarized beam splitter. A 10x microscope objective (0.25 NA) focused the beam on the sample. The Stokes-shifted backscattered photons were separated using a dichroic mirror (LP805, Thorlabs, Newton, NJ, USA) and coupled into a 0.22 NA multimode fiber (M105L02S-B, Thorlabs, Newton, NJ, USA) connected to the Raman spectrometer (HyperNova, StellarNet, USA) with a thermoelectrically cooled (-60°C) CCD camera (Andor Technology, Belfast, Northern Ireland).

Isolated EV samples were diluted 10^4^ times (see dilution response of SERS substrate in **Supplementary Figure 3B**) with distilled water and drop-casted on Au nanopillar-coated SERS substrates (Silmeco, Copenhagen, Denmark). A 1.3 µL drop of each sample was placed on a hydrophobic surface and dried for 15 minutes. The particles attached to gold nanopillars were measured with an exposure time of 1 second. The measurement region was scanned in 5 µm steps along 30 steps in each direction, resulting in 961 measurements per sample.

Due to the high dilution, some measurement sites did not contain extracellular vesicles (EVs) and only recorded surface responses. These blank measurements were filtered out during preprocessing using cluster analysis. The measurements were divided into five groups, and the groups whose signatures did not contain the predefined EV peaks were eliminated. After filtration, samples were calibrated using the polystyrene spectra taken before each measurement. The spectra were then smoothed and baseline corrected using Asymmetric Least Squares (ALS) algorithm. The baseline corrected spectra were normalized to the peak intensity around 642 cm^-1^ that is observed in most EV spectra.

### Statistical Analysis

#### in vitro Essay Data Analysis

Graphpad Prism 10 (Boston, MA) was used to analyze and graph data. All data is normalized to its corresponding well in the plate along the time course of the experiment. For each day of measurement, each well was also normalized to the average viability of the control wells. All data is presented as the mean ± standard deviation of the collected wells. IC_50_ of CCK8 and albumin graphs was determined by fitting the data to a non-linear regression model to create an inhibition dose response curve (concentration of inhibitor versus normalized response) with variable slope. Specifically, Graphpad Prism fit data to the following model: **Y=100/(1+(IC**_**50**_ **/X)^HillSlope)**, where Y is the normalized viability, X is the concentration of the inhibitor, in this case acetaminophen. IC_50_ and Hill Slope are calculated and output as a results of least squares regression.

#### Exploratory Analysis and Linear Regression on Raman Dataset

Principal Component Analysis (PCA) was employed to reduce the dimensionality of the Raman spectral data and identify patterns corresponding to different concentration levels. Preprocessed spectral data, normalized to minimize variations due to instrumental factors, served as input for PCA. The analysis focused on the first two principal components (PC1 and PC2), which explained the majority of the variance in the datasets. PCA was applied separately to data from Day 0 and Day 2 for both sets to assess differences in clustering over time. Scatter plots of PC1 versus PC2 were used to visualize the separation between concentration groups, with the percentage of variance explained by each component indicated in the plots. All PCA computations and visualizations were performed using the built-in functions in Matlab (MathWorks, Natick, MA, USA).

The baseline-corrected Raman spectra were used to construct a regression model utilizing Gaussian Process Regression (GPR) to predict concentration levels. The dataset comprised spectra for four different concentrations: 0 mM, 10 mM, 20 mM, and 40 mM. For each concentration group, balanced number of spectra were randomly selected to form the training set.

The spectra were split into predictor variables (intensity values) and response variables (concentration levels). The linear regression model was trained using the built-in *fitlm* function in MATLAB which performs the linear regression operation with several options. Predictions were made using the same dataset, and the RMSE was calculated. The process was iterated until convergence was achieved. The final RMSE and residuals were calculated and plotted, with predictions visualized using box plots to compare predicted concentrations against actual concentrations. All data visualizations were made using the built-in tools in Matlab. Error shading in MATLAB plots was performed using the *shadedErrorBar* function (Raoul Campell, available at https://github.com/raacampbell/shadedErrorBar, accessed Jan. 28, 2025). The exported plots were arranged and finalized in Inkscape (Version 1.3, available at https://inkscape.org). The writing and editing of this manuscript were supported by artificial intelligence (AI)-assisted tools to enhance clarity, coherence, and technical accuracy while ensuring compliance with journal guidelines

#### Correlation Matrix

The correlation matrix was calculated by determining the Pearson correlation coefficient between each pair of variables. This coefficient, *r*_*xy*_, is given by the equation:

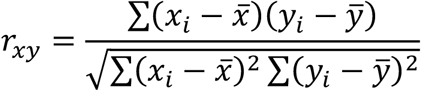

where *x*_*i*_ and *y*_*i*_ are individual data points, and 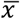 *and* 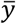 are the sample means. All calculations were performed using Matlab’s Statistics Toolbox.

## Acknowledgements

U.D. acknowledges the support of a seed award from Stanford University’s PhIND Center. We also acknowledge funding from the National Institutes of Health under project numbers R21GM140656 (L.B. and O.B.U.), R01AR081529 (A.G. and O.B.U.), and R21GM136002 (A.G. and O.B.U.). U.A. acknowledges funding from the European Union’s Horizon Europe research and innovation programme under the Marie Skłodowska-Curie grant agreement No. 101066038

## Author contributions

*Conceptualization:* U.P, D.A, O.B.U. and U.D.; *Methodology:* U.P, L.B, H.T., L.F., U.A., A.A.G., D.A. O.B.U. and U.D; *Software*: U.P and U.A; Data Curation, *Validation and Formal analysis:* U.P, L.B, H.T. and U.A.; *Investigation:* U.P, L.B, L.F., H.T., A.A.G. and U.A; *Writing-Original Draft*: U.P, L.B, U.A., D.A. O.B.U. and U.D; *Writing-Review and Editing*: U.P, L.B, H.T., L.F., U.A., A.A.G., D.A. O.B.U. and U.D; *Supervision, Project administration, Funding acquisition and Resources:* O.B.U. and U.D.

## Data availability

All data supporting the findings of this study are available within the paper and its Supplementary Information. The custom code used to generate Figures 4 and 5 has been deposited on Zenodo with the identifier https://doi.org/10.5281/zenodo.14768753.

Additional data that support the findings of this study are available from the corresponding author upon reasonable request.

