## Supplementary figure for "SHINE: SERS-based Hepatotoxicity detection using Inference from Nanoscale Extracellular vesicle content"

### Supporting Information

#### 1) Brightfield images from 12-well plate containing 2D rat hepatocyte cultures

**Supplementary Figure 1** shows representative brightfield micrographs of 2D rat hepatocytes cultured in Geltrex medium and exposed to increasing concentrations of APAP (0, 10, 20, and 40 mM). Images were collected one day (top two rows) and two days (bottom two rows) after the initial treatment. Each panel illustrates the dose-dependent morphological changes in hepatocytes, such as reduced cell density and altered cellular structure, which become more pronounced with higher APAP concentrations and longer exposure times. Scale bars (white) are provided in each micrograph to indicate the magnification.

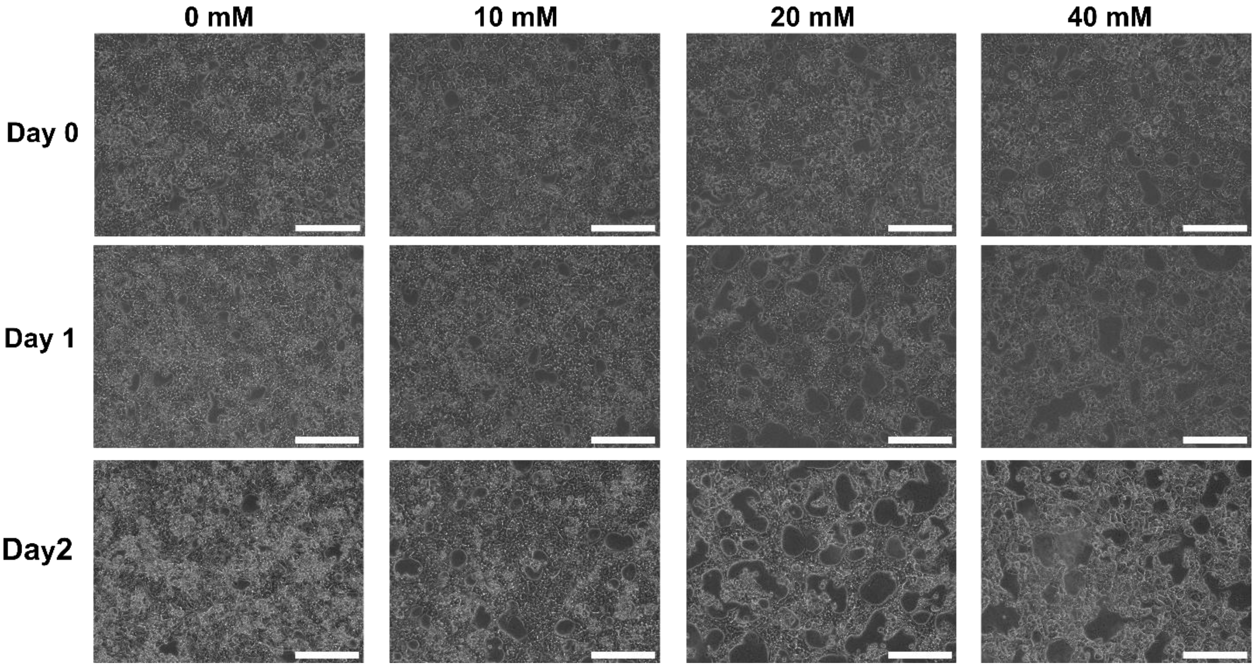


**Supplementary Figure 1:** Morphological changes of the primary rat hepatocytes cultured with Geltrex supplementation over two days of dosage with varying concentrations of Acetaminophen. Scale bar: 300 µm.

#### 2) PCA overlap quantification

Overlap between PCA clusters was calculated by Ellipsoid Overlap Volume Function: (EOVF). This method relies on calculating a metric called Bhattacharyya distance and then utilize these distances to find the overlap as shown in **Eq. S1** below:

$$Overlap = \exp\left(-Bhattacharyya\ distance\right) \qquad (Eq. S1)$$

Where Bhattacharyya distance is defined as,

$$D_B = \frac{1}{8}(\mu_1 - \mu_2)^T \Sigma^{-1}(\mu_1 - \mu_2) + \frac{1}{2}\ln\left(\frac{det\Sigma}{\sqrt{det\Sigma_1 \cdot det\Sigma_2}}\right) \qquad (Eq. S2)$$

This formula was applied pairwise on the PCA data and the overlap table in **Table S1** was obtained.

**Table S1:** Overlap between PCA clusters calculated by Ellipsoid Overlap Volume Function: (EOVF).

| Day 0 | Group 1 | Group 2 | Group 3 | Group 4 |
|---|---|---|---|---|
| **Group 1** | 1.000 | 0.5914 | 0.5067 | 0.7352 |
| **Group 2** | | 1.000 | 0.534 | 0.7679 |
| **Group 3** | | | 1.000 | 0.6448 |
| **Group 4** | | | | 1.000 |

| Day 2 | Group 1 | Group 2 | Group 3 | Group 4 |
|---|---|---|---|---|
| **Group 1** | 1.000 | 0.3359 | 0.2354 | 0.4902 |
| **Group 2** | | 1.000 | 0.271 | 0.5468 |
| **Group 3** | | | 1.000 | 0.3859 |
| **Group 4** | | | | 1.000 |

The rule to interpret the overlap evaluation is given in **Eq. S2**:

$$\left\{\begin{matrix} 0 < \mathrm{EOVF} < 0.4 & Low\ Overlap \\ 0.4 < \mathrm{EOVF} < 0.7 & Moderate\ Overlap \\ 0.7 < \mathrm{EOVF} < 1.0 & High\ Overlap \end{matrix}\right\} \qquad (Eq.S2)$$

This shows that the EOVF values between the class pairs are all in the low or moderate overlap regime for Day 2 measurements, while they are moderate or high for Day 0 samples.

#### 3) Size distributions of EVs obtained by NTA and iSCAT

We characterized EV sizes using iSCAT and found that the mean, median, and mode sizes showed slight variations between Day 0 and Day 2 across APAP doses (**Supplementary table 1**). At 40 mM, EV sizes were slightly larger compared to 0 mM, with mean sizes ranging from 115.2 nm to 122.9 nm.

**Supplementary table 1**: iSCAT size characterization. Mean, mode and median sizes of the EVs measured iSCAT. Units are nanometers.

| iSCAT | Day 0 | | Day 2 | |
|---|---|---|---|---|
| | **0 mM** | **40 mM** | **0 mM** | **40 mM** |
| **Mean (nm)** | 122.7 | 122.9 | 115.2 | 118 |
| **Median (nm)** | 118.5 | 118.9 | 100.3 | 113.4 |
| **Mode (nm)** | 137.2 | 138.1 | 87.3 | 100.5 |

We performed nanoparticle tracking analysis (NTA) to evaluate EV sizes and observed distinct differences between APAP doses and time points. The mean, mode, and median sizes, along with their standard errors, indicated that EV sizes tended to decrease from Day 0 to Day 2 at 40 mM, highlighting a dose- and time-dependent response as shown in **Supplementary table 2**.

**Supplementary table 2**: NTA size characterization. Mean, mode and median sizes of the EVs evaluated by nanoparticle tracking analysis. Standard errors were also given in the table.

| NTA | Day 0 | | Day 2 | |
|---|---|---|---|---|
| | **0 mM** | **40 mM** | **0 mM** | **40 mM** |
| **Mean (nm)** | 163.8±4.4 | 159.4±5.4 | 185.4±1.1 | 155.1±5.0 |
| **Mode (nm)** | 165.5±8.5 | 166.2±10.6 | 155.3±7.7 | 119.8±5.6 |
| **Median (nm)** | 161.5±5.8 | 169.2±7.0 | 132.5±16.9 | 131.8±2.5 |

#### 4) Flow cytometry additional data

We used flow cytometry to evaluate the presence of CD63+ extracellular vesicles (EVs) over time. Gating analysis showed that the EV population had high purity, with 99.7% and 99.9% of particles classified as EVs on Day 0 and Day 2, respectively (**Supplementary figure 2A, D**).

We observed a significant increase in CD63 expression within the EV population over time. On Day 0, only 0.76% of the gated EVs expressed the CD63 marker (**Supplementary figure 2B**), while this percentage increased sharply to 93.1% on Day 2 (**Supplementary figure 2E**).

We also analyzed the overlap of CD63+ markers with the EV population. On Day 0, CD63+ EVs accounted for just 2.23% of the total EV population (**Supplementary figure 2C**). By Day 2, this proportion increased dramatically to 93.4% (**Supplementary figure 2F**). These results demonstrate a clear time-dependent upregulation of CD63 expression in the EV population.

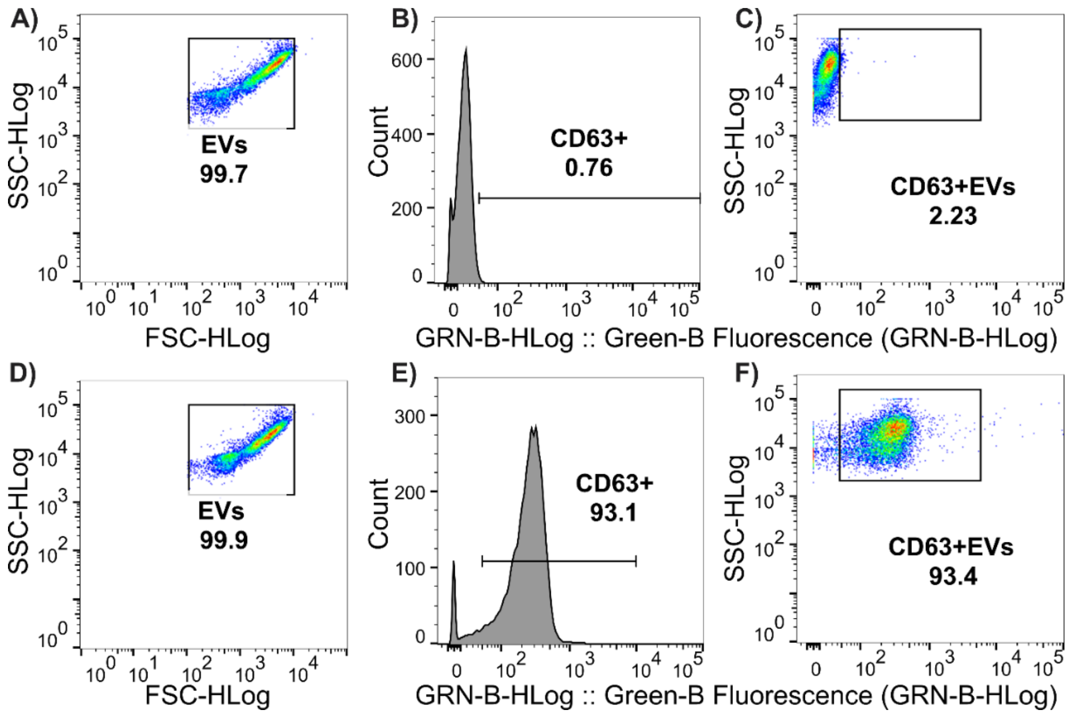


**Supplementary figure 2**: Flow cytometry forward scattering and side scattering analysis results. A-D) Forward scattering analysis for unstained (A) and CD63+ EVs. B-E) Fluorescence intensities vs number of particles detected in green channel. 2.24% of the unstained particles detected in the EV window. C-F) Side-scattered log intensities vs number of particles detected. 93.4% stained particles detected in the EV window.

### 5) SERS substrate data, dilution graph

We characterized the SERS response of the Au substrate and evaluated the dilution dependence of EV preparations. Supplementary figure 3A shows the spectral response map of the Au substrate before EV addition, highlighting uniform intensity across the surface at the 1576 cm⁻¹ wavenumber.

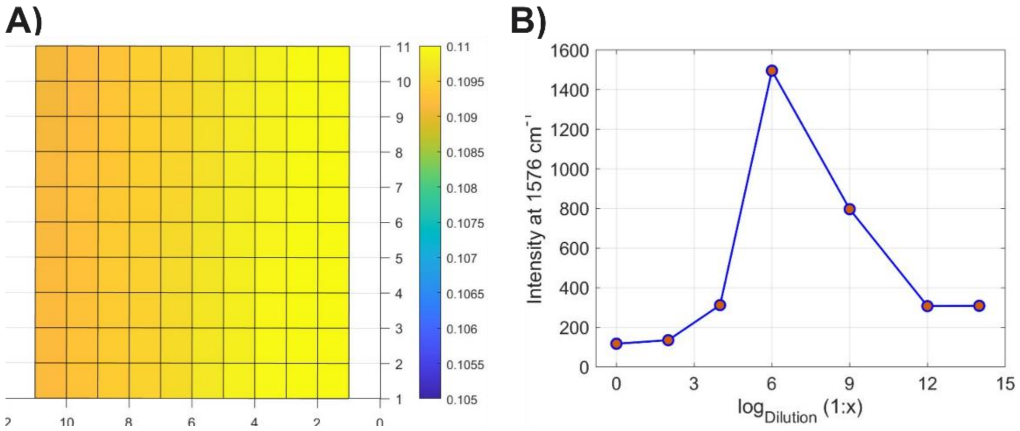


**Supplementary figure 3**: SERS characterization. A) Spectral response map of the Au substrate before EV addition.  B) Dilution dependence of the EV preparation on Au substrate.

In **Supplementary Figure 3B**, we investigated the dilution dependence of the EV preparation on the Au substrate by monitoring the SERS intensity at 1576 $cm^{-1}$. The results revealed a clear trend: the intensity increased with dilution, reaching a maximum at a $1:10^6$ dilution, followed by a decline at higher dilutions. This finding suggests an optimal concentration for EV binding and detection on the Au substrate. Since this test was performed using Day 0 samples exposed to 0 mM APAP, we adjusted the dilution across all samples to $1:10^4$ to ensure a sufficient number of EVs for analysis.
